# PANGEA: A New Gene Set Enrichment Tool for *Drosophila* and Common Research Organisms

**DOI:** 10.1101/2023.02.20.529262

**Authors:** Yanhui Hu, Aram Comjean, Helen Attrill, Giulia Antonazzo, Jim Thurmond, Fangge Li, Tiffany Chao, Stephanie E. Mohr, Nicholas H. Brown, Norbert Perrimon

## Abstract

Gene set enrichment analysis (GSEA) plays an important role in large-scale data analysis, helping scientists discover the underlying biological patterns over-represented in a gene list resulting from, for example, an ‘omics’ study. Gene Ontology (GO) annotation is the most frequently used classification mechanism for gene set definition. Here we present a new GSEA tool, PANGEA (PAthway, Network and Gene-set Enrichment Analysis; https://www.flyrnai.org/tools/pangea/), developed to allow a more flexible and configurable approach to data analysis using a variety of classification sets. PANGEA allows GO analysis to be performed on different sets of GO annotations, for example excluding high-throughput studies. Beyond GO, gene sets for pathway annotation and protein complex data from various resources as well as expression and disease annotation from the Alliance of Genome Resources (Alliance). In addition, visualisations of results are enhanced by providing an option to view network of gene set to gene relationships. The tool also allows comparison of multiple input gene lists and accompanying visualisation tools for quick and easy comparison. This new tool will facilitate GSEA for *Drosophila* and other major model organisms based on high-quality annotated information available for these species.

## INTRODUCTION

Modern genetics and genomics owe much to work done using common model organisms. These models continue to make a significant contribution to the understanding of development, metabolism, neuroscience, behaviour and disease. With the onset of the ‘big data’ era has come a need for analysis platforms that deconvolute complex data from multispecies studies. Model organism databases (MODs) are knowledgebases dedicated to the curation, storage and integration of species-specific data for their research community. The past decade has seen a number of efforts aimed at pulling together model organism and human data to facilitate a more interdisciplinary approach; examples include MARRVEL (Model organism Aggregated Resources for Rare Variant ExpLoration) (1), Gene2Function (2) and the Monarch Initiative (3). Furthermore, the Alliance of Genome Resources (4), a consortium of seven model organism databases and the Gene Ontology (GO) Consortium (GOC), formed recently with the objective of building an umbrella resource from which users can navigate combined data within a single integrated knowledgebase. To help support such a resource, the MODs are working to reduce the divergence in the way the primary data is curated, stored and presented to facilitate comparative and translational research.

Although these integrated resources allow search and comparison across certain data classes, large-scale data analysis remains in the domain of stand-alone bioinformatic tools such as DAVID (5), TermMapper (6), GOrilla (7), PANTHER Gene List Analysis (8), WebGestalt (9) and g:Profiler (10), with a focus on processing gene list to extract a statistical measure of shared biological features, usually termed gene set enrichment analysis (GSEA). The most frequently used gene set classification in GSEA is GO annotation, which is based on the most widely-used ontology (a hierarchical controlled vocabulary) in biological research for the wild-type molecular function(s), biological process(es) and cellular component(s) associated with a given gene product (11,12). Many GSEA tools also incorporate classifications from other sources such as Reactome (13) and KEGG (14) pathways (e.g., DAVID, WebGestalt and g:Profiler) and, for human studies, there may be additional data sources such as the Human Phenotype Ontology (HPO) (15) and Online Mendelian Inheritance in Man (OMIM) (16) (e.g., WebGestalt). The addition of gene sets beyond the GO allows users to extract more classification information, as well as seek trends and overlap in the enriched sets. DAVID and g:Profiler are two of the few resources that make it possible for users to compare different sets on the same display; however, the ability to interact with the results post-processing at these resources is rather limited.

Despite the abundance of tools, we found that they do not fully meet community needs, primarily because they were overly focused on human gene data and were not using the most up to date data on other species. For example, Reactome pathway annotation is based on computational predictions derived from manually curated human pathways. There are a few organism-targeted analysis tools; the prokaryote-centred GSEA FUNAGE-Pro (17) is one example in which the underlying knowledgebase was assembled to cater to the needs of a specific research community. A number of useful gene classification resources (e.g. pathways, complexes, gene groups) have been developed by the Drosophila RNAi Screening Center (DRSC) and FlyBase, the Drosophila knowledgebase (18–22). Furthermore, in FlyBase and indeed across the MODs, there are several types of curated data in common, including disease models, phenotypes and gene expression, which could be used for GSEA. In contrast to the annotation of gene function data with GO, which is done in a consistent manner across multiple organisms, other data types are represented within the MODs in diverse ways, reflecting some of the technical differences in the genetics of these organisms. The Alliance was founded to integrate data across many MODs (4) and now provides a source of harmonised data that can also be used for GSEA. To take full advantage of the research in diverse model organisms, we describe our creation of a new tool that we name PANGEA. Although our primary focus was on Drosophila genes, we developed PANGEA to also include rat, mouse, zebrafish, nematode worm data, as well as harmonised human data to facilitate translational research. PANGEA not only incorporates additional gene set classifications from Alliance and MODs, but also have implemented the features that enhance the presentation of enrichment results by allowing the user to select sets and compare them visually to facilitate interpretation as well as making it easy to do parallel GSEA for multiple gene lists. This flexibility enables users to adapt the tool to their needs and allow ‘fortuitous’ discovery by widening the pool of knowledge for the purpose of analysis.

## MATERIAL AND METHODS

### Building the knowledgebase for PANGEA

The gene set classification is a way to group genes based on commonality such as the same biological pathway. We have collected more than 300,000 gene sets from various public resources (Table 1,2) for fruit fly *D. melanogaster*, the nematode worm *C. elegans*, the zebrafish *D. rerio*, the mouse *M. musculus*, the rat *R. norvegicus* and human *H. sapiens*. For annotations based on a controlled vocabulary arranged in hierarchical structure, such as gene group and phenotype annotations from FlyBase, gene-to-gene set relationships were assembled after the hierarchical structure was flattened. An exception was made for GO annotations, which were assembled in two ways, with and without being flattened, allowing users to choose which output is used in the analysis. GO annotations include evidence codes that indicate the type of evidence supporting the annotation. For example, ‘IDA’ means that an annotation was supported by a direct assay, whereas ‘ISS’ means that the annotation was inferred from sequence or structural similarity. Using such evidence codes, GO gene sets were built with additional configurations: 1.) subsets based on experimental evidence codes, i.e., excluding annotations only based on phylogenetic, sequence or structural similarity and other computational analyses (IEA, IBA, IBD, IKR, IRD, ISS, ISO, ISA, ISM, IGC, RCA); 2.) subsets excluding annotations only supported by high-throughput (HTP) evidence codes (HTP, HDA, HMP, HGI and HEP); 3.) subsets of GO generic terms (GO slim) provided by the GOC (http://geneontology.org/docs/download-ontology/#subsets); 4.) subsets of very high-level GO term classifications used by FlyBase and the Alliance originally generated to support GO summary ribbon displays. For Drosophila phenotype annotations from FlyBase, we assembled the gene-to-phenotype association using the “genotype_phenotype_data” file available in the FlyBase Downloads page, in which phenotypes are associated with individual genotypes and controlled vocabulary identifiers are indicated. This allowed us to extract only those genotypes where we could be certain that the phenotype was associated with the perturbation of a single Drosophila melanogaster gene (i.e. single classical or insertional alleles). Because different resources use different gene or protein identifiers, we used an inhouse mapping program to synchronise IDs to NCBI Entrez gene IDs, official gene symbols and gene identifiers of species-specific resources, such as MGI and FlyBase (Table 1). The PANGEA knowledgebase stores the information of gene set classification, gene annotation obtained from NCBI as well as the information for ID mapping among various resources.

**Table 1.**
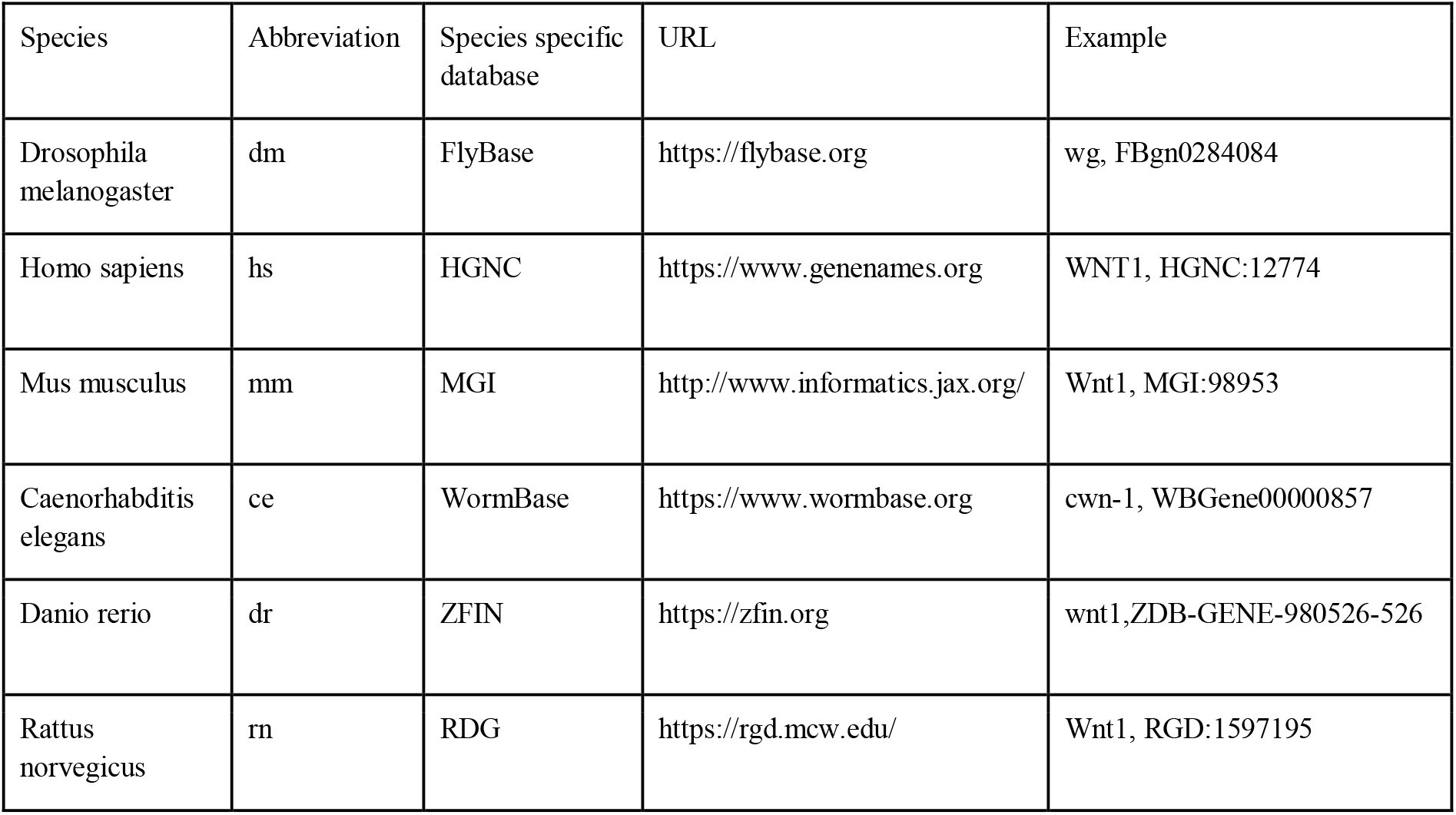
Species coverage by PANGEA and corresponding species-specific databases.

**Table 2.** Knowledgebase of PANGEA built from various gene annotation resources.

| Type | Source | URL | Species covered at PANGEA |
| --- | --- | --- | --- |
| Gene Ontology | GO | http://geneontology.org/ | hs,mm,rn,dr,dm,ce |
| pathway | KEGG | https://www.genome.jp/kegg/ | hs,mm,rn,dr,dm,ce |
| pathway | Reactome | https://reactome.org/ | hs,mm,rn,dr,dm,ce |
| pathway | PantherDB | http://www.pantherdb.org/ | dm |
| pathway | FlyBase | https://flybase.org/ | dm |
| pathway | PathON | https://www.flyrnai.org/tools/pathon/ | dm |
| group | HGNC | https://www.genenames.org/ | hs |
| group | FlyBase | https://flybase.org/ | dm |
| group | GLAD | https://www.flyrnai.org/tools/glad/ | dm |
| protein | COMPLEAT | https://www.flyrnai.org/compleat/ | dm |
| protein | EBI complex portal | https://www.ebi.ac.uk/complexportal | hs,mm,rn,dr,dm,ce |
| phenotype | Alliance of genome resources | https://www.alliancegenome.org/ | hs,mm,rn,dr,dm,ce |
| phenotype | FlyBase phenotype | https://flybase.org/ | dm |
| expression | Alliance of genome resources | https://www.alliancegenome.org/ | mm,rn,dr,dm,ce |

### Building gene sets of preferred tissue expression

To study the diversity and dynamics of the Drosophila transcriptome, the modENCODE consortium sequenced the transcriptome in twenty-nine dissected tissues (23) and the processed datasets are available at FlyBase (http://ftp.flybase.net/releases/current/precomputed_files/genes/). A program was implemented in the Python programming language to identify genes expressed at a substantially higher levels in one tissue versus any other tissue. This program first groups the RNA-seq datasets based on tissue. For example, all data related to the nervous system are grouped together. It then calculates the average reads per kilobase per million mapped reads (RPKM) expression values for each gene in each tissue group. Genes were identified as preferentially expressed in a given tissue group if their average expression in the tissue group is three-fold or higher than the average expression in any other tissue group. Genes with average RPKM value lower than 10 were excluded. Genes defined in this way as ‘tissue-specific’ then get annotated with the relevant tissue to generate the tissues expression classification set.

### Datasets used for testing

Drosophila cell RNAi screen phenotype data was obtained from the DRSC (https://www.flyrnai.org/) via download of a file of all available public screen ‘hits’ (results) (https://www.flyrnai.org/RNAi_all_hits.txt). RNAi reagents of optimal design were selected. The criteria for optimal design were no CAN or CAR repeat, fewer than 6 predicted OTEs (off-target alignment sites of 19 bp) and a single gene target. RNAi reagents were mapped to current FlyBase gene identifiers using a DRSC internal mapping tool. Screens focused on major signalling pathways were selected for PANGEA analysis (24–29). Proteomics data was obtained from Tang *et al*., (2021) (30) and high-confident prey proteins identified by mass-spec (Supplementary Table 2) were used for analysis.

### Gene set enrichment statistics used at PANGEA

Hypergeometric testing is performed to calculate P values for GSEA using the PypeR function in R. Bonferroni correction for multiple statistical tests, Benjamini-Hochberg procedure for false discovery rate adjustment, and Benjamini-Yekutieli procedure for false discovery rate adjustment were performed using the p.adjust function in R.

### Web tool implementation

PANGEA is a SaaS (Software as a Service) web tool (https://www.flyrnai.org/tools/pangea/) and is built following a three-tier model, with a web-based user-interface at the front end, the knowledgebase at the backend, and the business logic in the middle tier communicating between the front and back ends by matching input genes with gene sets, doing statistical analysis and building visualization graphs. The front page is written in PHP using the Symfony framework and front-end HTML pages using the Twig template engine. The JQuery JavaScript library is used to facilitate Ajax calls to the back end, with the DataTables plugin for displaying table views and Cytoscape and VegaLite packages for the data visualisations. The Bootstrap framework and some custom CSS are used on the user interface. A mySQL database is used to store the knowledgebase. Both the website and databases are hosted on the O2 high-performance computing cluster, which is made available by the Research Computing group at Harvard Medical School.

## RESULTS

### Preparation of the classified gene sets for GSEA: the PANGEA knowledgebase

GSEA relies on high-quality annotation of genes/gene products with information related to their biological functions. For PANGEA, we used multiple sources of annotation to generate more than 300,000 different classes of gene function for five major model organisms (*D. melanogaster, C. elegans, D. rerio, M. musculus, R. norvegicus*) and human. For example, pathway annotations allow users to identify metabolic or signalling pathways that are over-represented in a gene list and help understand causal mechanisms underlying the observed phenotype from a screen. Pathway annotations from KEGG, PantherDB, and Reactome, as well as manually curated Drosophila gene sets, such as FlyBase Signalling Pathways and the DRSC PathON annotation (18,21), are included in the PANGEA knowledgebase.

The GO annotation set provides the_comprehensive knowledge on gene functions and we store the gene-to-gene set relationships from GO in two ways. One is the direct gene-to-GO term associations as obtained from the gene association file while the other stores the gene-to-GO term associations with consideration given to child-parent relationships. The latter is recommended for use in GSEA as it reflects the intended use of the ontology in curation practice. The direct, gene-to-term set may be useful to understand the depth of annotation for each gene. In addition, we also generated two gene annotation subsets using evidence codes. The “experimental data only” subset includes only those gene associations that are supported by experimental evidence codes. The “excluding high-throughput experiments’’ subset excludes annotations only supported by HTP evidence codes. Excluding HTP data may be important to avoid bias when analysing similar studies (31). GO slim subsets are the cut-down versions of GO that give a broad overview of the ontology content without the detail of the specific, fine-grained terms. The PANGEA knowledgebase includes two sets of GO slim annotations from different resources.

In addition to GO and pathway annotations, MODs provide organism-specific curation of important aspects of gene information, such as gene expression and mutant phenotype, that are not captured in GO. The Alliance is focused on the harmonisation and centralisation of major MODs data (4,32). To take advantage of this effort, we integrated gene-to-tissue expression and gene-to-disease (model) association annotations from the Alliance into the PANGEA knowledgebase. As all organisms in the Alliance use the Disease Ontology (DO) for annotation, this set is easily comparable across species. The Alliance DO annotation set also includes disease association to model organism genes via an electronic pipeline using orthology with human disease genes which expands the set provide by the MODs. Moreover, for Drosophila genes we assembled an additional gene set from phenotype annotation at FlyBase by extracting phenotype data associated with a ‘single allele’ genotype (i.e., single classical or insertional alleles), allowing users to perform meaningful enrichment analyses on this data class for the first time.

Also included in PANGEA are gene group classification (eg. kinases and transcription factors) from organism-specific resources (human and fly), protein complex annotations for multiple organisms from the EMBL-EBI Complex Portal (33) and COMPLEAT (22), and bespoke gene sets using Drosophila modENcode RNAseq data to identify genes particularly highly expressed in one tissue (see Materials and Methods).

In summary, we have assembled more than 300,000 different gene sets that can be used in PANGEA to assess the enrichment of particular biological features in an input gene list.

### Features of the PANGEA user interface

GSEA can be computationally intensive because of the number of gene sets being tested and potentially large number of genes entered by users. Therefore, the step of pre-processing user’s input by mapping the input gene identifiers to the gene identifiers used for gene set annotation, is set up as a standalone ID mapping page (accessed by clicking ‘Gene Id Mapping’ on the top toolbar) instead of combining it with the analysis step. Gene identifiers supported by PANGEA include Entrez Gene IDs, official gene symbols and primary gene identifiers from MODs. Users might need to analyse lists of other identifiers such as UniProtKB IDs and Ensembl gene IDs. Users can use “Gene Id Mapping” tool and select an organism of choice to map IDs. As gene annotation is an on-going process, the gene identifiers as well as gene symbols might change over time. Even with the same type of gene identifiers such as FlyBase gene ID, the IDs used by users might be from a different FlyBase release. Therefore, ID-mapping step is an optional but recommended first step to ensure that the entered IDs are synchronized with the IDs used by PANGEA gene set annotation. Users of FlyBase may also directly export a ‘HitList’ of genes generated in FlyBase to the tool by selecting the ‘PANGEA Enrichment Tool (DRSC)’ from the dropdown ‘Export’ menu. An option for users to upload a background gene list for the analysis is provided; this may be useful when analysing hits from a focused screen using a kinase sub-library instead of a genome-scale library, for example. PANGEA identifies all relevant gene sets and provides enrichment statistics such as p-values, adjusted p-values, and fold enrichment, as well as the genes shared by the input gene list and gene set members. Users have the option to set different p-value cut-offs and visualise the results using a bar graph, the height and colour intensity of which can be customised (Figure 1A). In addition, users can select gene sets of interest to examine the overlap of genes in different gene sets using the ‘Gene Set Node Graph’ visualisation option. Nodes of different shapes in the network indicate genes or gene sets while edges reflect the gene-to-gene set relationship. This type of visualisation can help users identify the most relevant genes in each gene set as well as commonly shared or distinct gene members of the selected gene sets (Figure 1B,1C).

**Figure 1:**
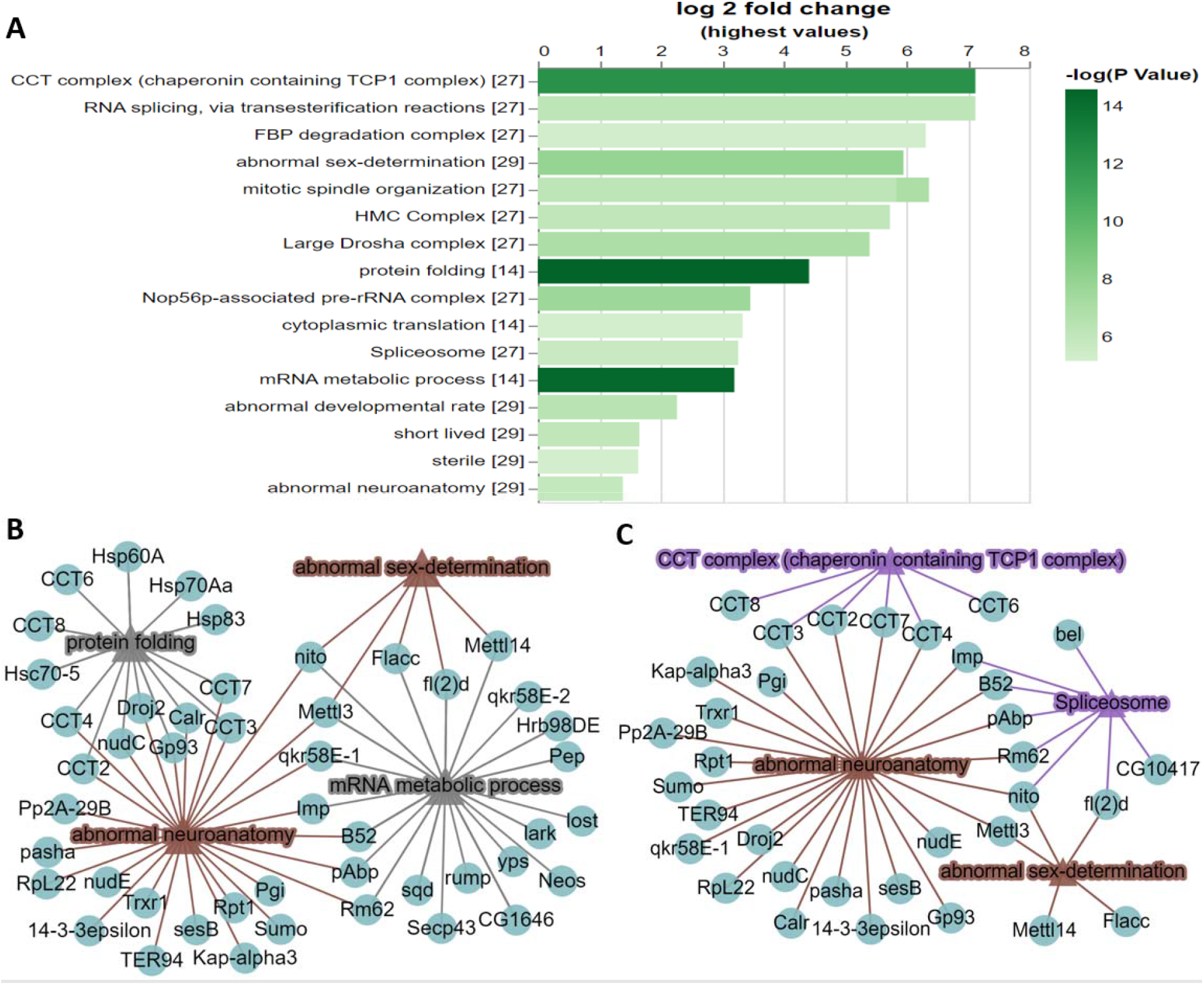
Example of analysing a single gene list using PANGEA. A proteomic interaction dataset was selected from a study of the m6A methyltransferase complex MTC (30). The 75 high-confidence interactors of the 4 subunits of the MTC (METTL3, METTL14, Fl(2)d and Nito) identified by affinity-purified mass spectrometry from *Drosophila* S2R+ cells were submitted via the ‘Search Single’ option at PANGEA and enrichment analysis was performed over phenotype, GO SLIM2 BP and protein complex annotation from COMPLEAT (literature based). The result was filtered using P value 1×10^-5^ cut-off and was illustrated as A) a bar graph and B) a network graph of selected gene sets from phenotype annotation and GO SLIM2 BP. Triangle nodes represent gene sets and circle nodes represent genes while the edges represent gene to gene set associations. C.) a network graph for selected gene sets from phenotype annotation and COMPLEAT protein complex annotation (literature based).

An under-appreciated use of GSEA tools is that researchers often use them as simple gene classification tools, for example, asking “which genes in my list are kinases?” to help inform further computational or experimental analysis. Having diverse classification sets is important because depending on the type of data/experiment being analysed, different gene sets may be more useful than others. It is often useful to be able to compare similar gene sets from different sources to help evaluate the evidence for support. In addition, PANGEA not only reports genes in an enriched gene set but also reports genes not covered by the gene set category selected, which may be interesting because of their lack of characterisation. This feature can help user answer question like “which genes in my list are not covered by KEGG annotation?”.

Often users need to analyse multiple gene lists and compare the results; however, majority of current web-based tools only allow the analysis of a single input list (plus background). Thus, users have to perform comparisons manually or using different tools. To address this need, PANGEA allows users to input multiple gene lists and compare results directly via a heatmap or a dot plot visualisation. For example, users might input gene hits from different phenotypic screens and compare what pathways, gene groups or biological processes that are common or different among results (Figure 2).

**Figure 2:**
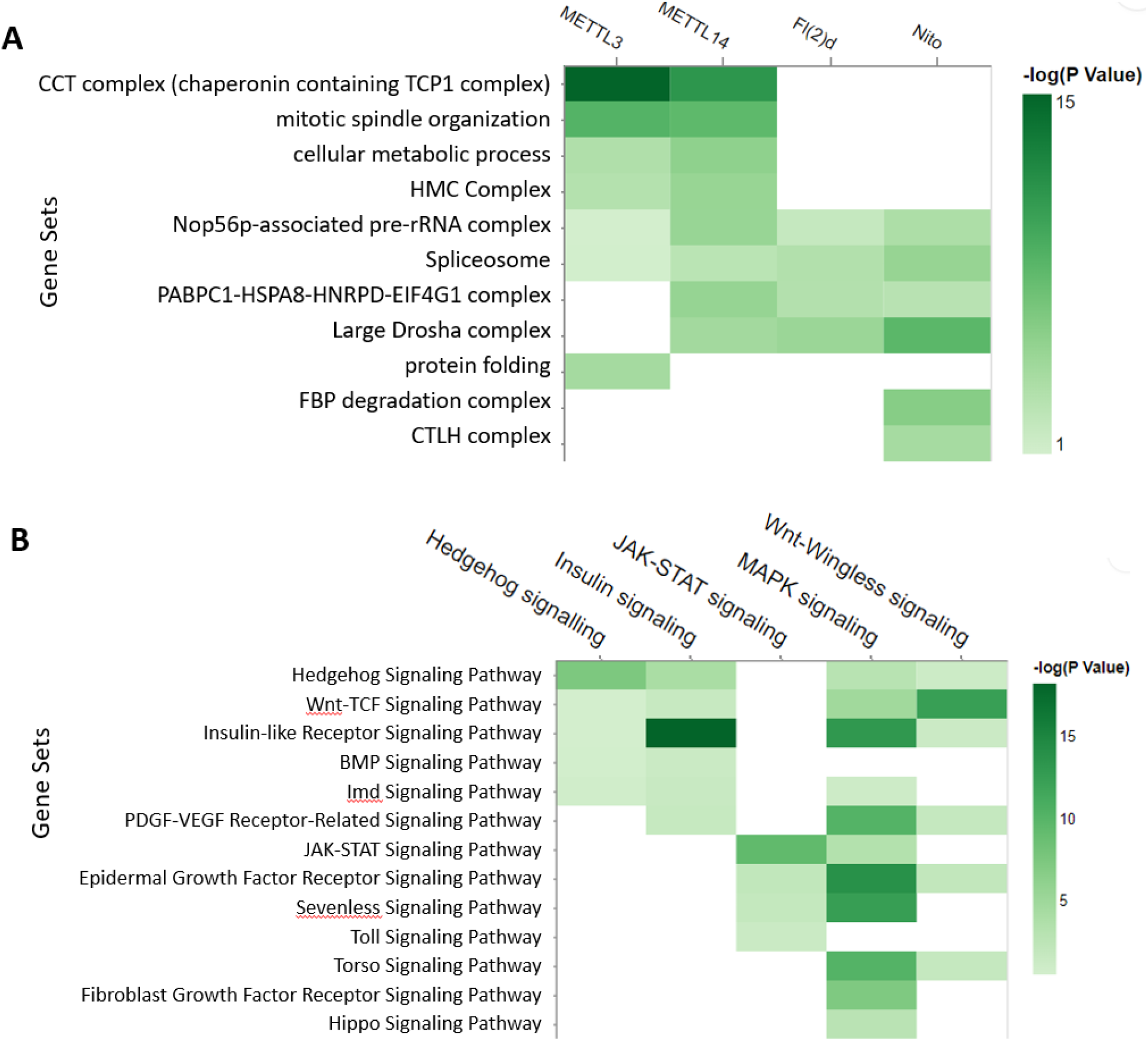
Examples of analysing multiple gene lists using PANGEA. A.) The prey proteins of multiple baits from AP mass-spec dataset (30) were submitted via the ‘Search Multiple’ option at PANGEA. The gene sets of protein complex annotation from COMPLEAT were selected. The comparison of enrichment over annotated protein complexes from the interacting proteins of four different baits was illustrated using a heatmap. B.) RNAi screen data of signalling pathway studies were obtained from DRSC RNAi data repository (24) and the hits were submitted via the ‘Search Multiple’ option at PANGEA. The gene sets of FlyBase pathway annotation were used. The comparison of the enrichment of signalling pathway components from the screen hits of five studies was illustrated using a heatmap.

### Testing the utility of PANGEA

To test the utility of PANGEA, we first analysed a proteomic interaction dataset from a study of the m6A methyltransferase complex MTC (30). In this study, using individual pull-downs of the four subunits (METTL3, METTL14, Fl(2)d and Nito) of the MTC complex, high-confidence interactors were identified by mass-spectrometry from *Drosophila* S2R+ cells. Submitting the combined list of 75 interacting proteins from the four baits via the ‘Search Single’ option at PANGEA (accessed by clicking ‘Search Single’ on the top toolbar) for Fly and performing GSEA over phenotype, SLIM2 GO BP as well as protein complex annotation from COMPLEAT (literature based) enrichment, we identified mRNA metabolic process (GO:0016071), protein folding (GO:0006457), abnormal sex-determination (FBcv:0000436), abnormal neuroanatomy (FBcv:0000435), CCT complex and Spliceosome complex among the top enriched gene sets with the most significant p-values (all <1×10^-5^) (Figure 1A). Next, we visualised SLIM2 GO BP and phenotype annotations using a network graph. GO mRNA metabolic process hits overlap with both abnormal sex-determination and abnormal neuroanatomy phenotypes, but GO protein folding hits only overlap with the abnormal neuroanatomy phenotype (Figure 1B). We also visualised protein complex and phenotype annotations using a different network graph, showing Spliceosome hits overlap with the phenotypes of abnormal sex-determination and abnormal neuroanatomy while the CCT complex hits only overlap with the abnormal neuroanatomy phenotype (Figure 1C). Enrichment of sex/reproductive phenotypes align with known function of MTC in regulating the splicing of female-specific Sex lethal (Sxl) and its roles in alternative splicing and sexual dimorphism, as well as the germ stem cell differentiation in the ovary (34). These GSEA results are also concordant with the fact that MTC is also known to have a significant role in neuronal mRNA regulation. The benefit of the network visualization is apparent when viewing how the gene set assignment overlap (Figure 1B,1C), which reveals that some of the MTC interacting proteins are associated with abnormal neuroanatomy phenotype and that the mechanism of the association is through the CCT complex in the process of protein folding. In contract, the interacting proteins from Spliceosome have more broad impacts related to both abnormal neuroanatomy phenotype and abnormal sex-determination phenotypes through mRNA metabolic process.

We further analysed the protein complexes associated with each individual subunit using the ‘Search multiple’ option at PANGEA (accessed by clicking ‘Search Multiple’ on the top toolbar) and inputting the interacting protein lists for each bait, then compared the enrichment result using a heatmap visualisation (Figure 2A). The results indicated that some complexes, such as spliceosome subunits, are common to all

MTC subunits, whereas some are more specific, such as protein complexes CCT complex for METTL14 and METTL13. In addition, we further analysed phenotype enrichment for proteins associated with each individual subunit using the ‘Search multiple’ option, and comparison of the enrichment results shows many overlapping phenotypes, particularly with regard to sterility (Supplementary figure 1).

At another use case, we looked at phenotypic cell screen data. Large-scale RNA interference (RNAi) screening is a powerful method for functional studies in *Drosophila*. At the DRSC, datasets generated from more than one hundred screens are publicly available (24). We selected five screens designed to identify the genes for major signalling pathways and performed a GSEA analysis of the hits using the multiple gene list enrichment function of PANGEA. FlyBase signalling pathway gene sets were selected and the results of the five screens were compared side-by-side using a heatmap, which clearly illustrated enrichment of the core components of the corresponding pathways, as well as potential cross-talks between pathways (Figure 2B).

These use cases of PANGEA for phenotype screening data as well as proteomics data demonstrate the value of the tool in validating screen results as well as generating new hypotheses for further study.

## DISCUSSION

GSEA is a computational method used to identify significantly over-represented gene classes within an input gene list(s) by testing against gene sets assembled based on prior knowledge. Input gene lists are typically from high-throughput screens or analyses. Here we present PANGEA, a newly developed GSEA tool with major model organisms as its focus, that includes gene sets that are usually not utilized by other GSEA tools, such as expression and disease annotations from the Alliance, phenotype annotations from FlyBase, and GO subsets with different configurations. PANGEA is easy to use and has new features such as allowing enrichment analyses for multiple input gene lists and generating graphical outputs that make comparisons straightforward for users. In addition to the use cases presented here, i.e., analysing phenotypic screening and proteomic data, we anticipate that the tool will also facilitate analysis of gene lists from other types of data. For example, analysis of single-cell RNA-seq datasets at PANGEA might help users identify pathways and biological processes that are characteristic of various cell types. Users will also be able to answer questions on classification such as, “which genes in this list are kinases?”. PANGEA is designed to accommodate a wide range of biological data types and questions, providing users with a web-based analysis tool that is easily accessible and user-friendly.

We also note that gene classifications are not static, and the generic design of the tool means that it will be easy to update or expand PANGEA for more gene set classification and/or more species. In developing PANGEA, we sought to improve the effectiveness of GSEA by 1) providing multiple collections of genes classified by their function in different ways (classified gene sets); 2) ensuring the data underlying the classification of gene function was up to date; and 3) improving the visualization so that results from multiple gene sets or multiple gene lists could be compared easily.

## AVAILABILITY

The online resource is available without restriction at https://www.flyrnai.org/tools/pangea/.

## SUPPLEMENTARY DATA

Supplementary Data are available at NAR online.

## ACKNOWLEDGEMENT

We would like to thank the members of the Perrimon laboratory, the FlyBase consortium, the Drosophila RNAi Screening Center (DRSC), and the Transgenic RNAi Project (TRiP) for the discussion and suggestions during the design and implementation of the tool as well as the feedback during the tool testing. Additional thanks to Gil dos Santos (Harvard, US) and Gillian Millburn (Cambridge, UK) at FlyBase for their genotype-to-phenotype work.

