## supplementary figure 1 for "PANGEA: A New Gene Set Enrichment Tool for *Drosophila* and Common Research Organisms"

Supplementary Figure 1: The proteomic dataset of  high-confident preys of four bait proteins ( Fl(2)d, METTL14, METTL3, Nito) was obtained from the supplementary table 2 of the published study by Tang et al (PMID: 33649236, https://www.ncbi.nlm.nih.gov/pmc/articles/PMC7958400/). PANGEA "Search Multiple" function was used and Phenotype annotation was selected. The comparison of the enrichment of *Drosophila* phenotype annotation from the interacting proteins of four different baits was illustrated using a heatmap.

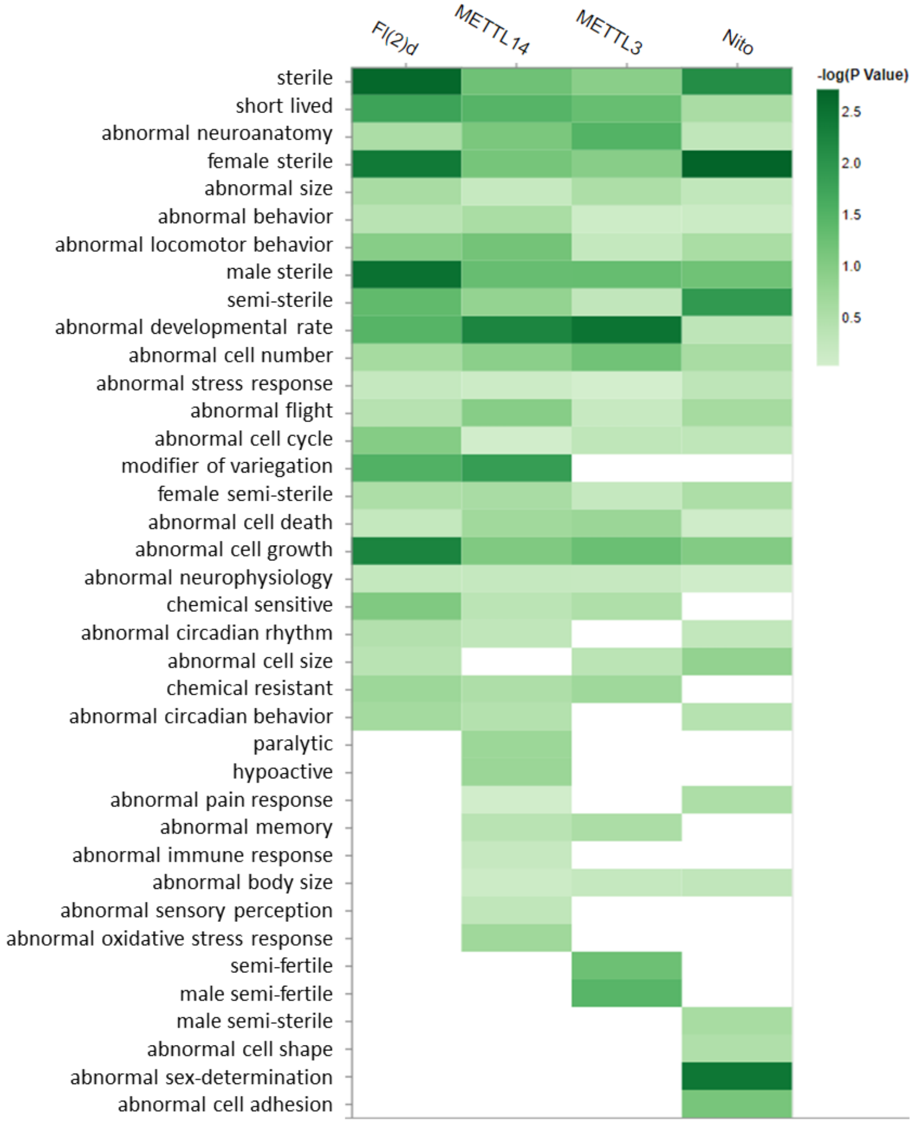
